# Ranking the direct threats to biodiversity in sub-Saharan Africa

**DOI:** 10.1101/2020.07.22.175513

**Authors:** Craig Leisher, Nathaniel Robinson, Matthew Brown, Deo Kujirakwinja, Mauricio Castro Schmitz, Michelle Wieland, David Wilkie

## Abstract

Sub-Saharan Africa benefits from large investments in biodiversity conservation, yet there is no prioritization of the many direct threats to biodiversity available to inform organizations developing sub-Saharan or sub-regional conservation strategies. Consequently, regional investments by funders of biodiversity conservation such as international conservation organizations, foundations, and bilateral and multilateral donors may be suboptimal. To identify the priority threats to biodiversity in sub-Saharan Africa, we classified the direct threats to biodiversity using standardized threats categories and triangulated data from a Delphi consensus of sub-Saharan Africa biodiversity experts, known threats to IUCN Red-listed sub-Saharan African species, and National Biodiversity Strategy and Action Plans from 47 sub-Saharan African countries. After ranking the threats from each source and averaging the rankings, we find that the highest threats are: annual and perennial crops (non-timber); logging and wood harvesting (natural forests); fishing and harvesting aquatic resources (marine and freshwater); and hunting and collecting terrestrial animals. Within the sub-regions of sub-Saharan Africa there is considerable variation. The highest ranked threat in Central Africa is hunting. In East Africa, it is agriculture. In Southern Africa, it is invasive non-native/alien species, and in West Africa, agriculture and logging are tied as the highest threats. There are known ways to address all of these threats, and concentrating investments on these threats while accounting for unique socio-ecological contexts across sub-Saharan Africa is essential for the sustained conservation of biodiversity.

## 1 Introduction

Conservation is funding constrained (Bruner et al. 2004; McCarthy et al. 2012; Wilkie et al. 2001), and suboptimal prioritization of limited resources reduces the benefits of conservation investments (McDonald-Madden et al. 2010). A number of global conservation prioritization assessments help guide limited resources to areas of high conservation value (Brooks et al. 2006; Hoekstra et al. 2010; Mittermeier et al. 1998; Myers et al. 2000; Olson and Dinerstein 1998; Olson et al. 2001; Stattersfield et al. 1998; UNEP 2019; Venter et al. 2016). These assessments highlight priority areas for conservation, but they do not specify the threats to biodiversity within the priority areas. While there are global assessments of biodiversity threats (Díaz et al. 2019; Dudgeon et al. 2006; Maxwell et al. 2016), there are known data gaps in understanding the top threats to biodiversity in many regions including sub-Saharan Africa (Joppa et al. 2016).

Sub-Saharan Africa is a global biodiversity conservation priority (Brooks et al. 2006) and includes irreplaceable bird and biodiversity areas (Di Marco et al. 2016), the iconic savannas of East Africa (Reid 2012), and one of the most biodiverse landscapes in the world, the Cape Floristic Province (Goldblatt 1997). Sub-Saharan Africa contains “crisis” and “very high risk” ecoregions with severe habitat conversion and minimal protected area coverage (Watson et al. 2016). Sub-Saharan Africa is also one of the poorest regions of the world, and 27 of the 31 countries classified as “low income” by the World Bank are in sub-Saharan Africa (World Bank 2019). These economic dynamics coupled with its high global biodiversity significance are two reasons why sub-Saharan Africa received in recent years more Official Development Assistance (ODA) for biodiversity conservation than any other region globally (Table 1).

**Table 1.** Biodiversity conservation Official Development Assistance gross disbursements, 2008-17 (US$ millions, 2017 constant prices)

| Region | Biodiversity ODA |
| --- | --- |
| Sub-Saharan Africa | \$2,068 |
| Asia | \$1,829 |
| South America | \$1,370 |
| North America and Central America | \$993 |
| Europe | \$290 |
| Oceania | \$69 |

Source: OECD-CRS (2019)

Funders of biodiversity conservation often decide on investments based on multi-year strategies driven by a set of priorities. Evidence-informed decision making (Pressey et al. 2017) to establish these priorities could improve the outcomes of conservation investments. The absence of a systematic and comprehensive assessments of the most prevalent biodiversity threats in sub-Saharan Africa hampers evidence-informed decision making by major donors who fund conservation across sub-Saharan Africa.

It also hampers systematic conservation planning at the local level (Margules and Pressey 2000). The lack of regional and sub-regional threat assessments reduces the efficacy of conservation planning at the local level. Knowing the regional and sub-regional direct threats helps local conservation projects better understand if addressing a threat locally would be sufficient to reduce or mitigate the threat given that local threat can be driven by macro forces that require a larger scope than an local conservation project. Knowing the regional and sub-regional threats also helps local conservation practitioners identify where partnerships and coalitions are needed to address specific threats at a scale meaningful to conservation.

There are several existing threat assessments that cover part or all of sub-Saharan Africa. The Intergovernmental Science-Policy Platform on Biodiversity and Ecosystem Services’ (IPBES) regional assessment report for Africa is a detailed assessment of the how the decline and loss of biodiversity is reducing nature’s contributions to people in Africa and how transformational development pathways based on ‘green’ and ‘blue’ economies can enable human well-being improvements that do not come at the expense of the environment (IPBES 2018). This report includes a qualitative assessment of the drivers of biodiversity change, but given its focus on exploring options for policymakers, the IPBES report does not attempt to prioritize direct threats to biodiversity. BirdLife’s Important Bird and Biodiversity Areas (IBAs) database includes threat (a.k.a. “pressure”) assessments for many IBAs in Africa, and two Africa-wide assessments of IBAs included an assessment of threats (BirdLife International 2011; Buchanan et al. 2009). IBA threat assessments are aggregated into an overall “impact score of threat” per site ranging from 0 to 9 (BirdLife International 2006). This threat score is useful for prioritizing IBAs based on threat levels, but it is difficult to categorize and summarize threats using IBA data. National Biodiversity Strategy and Action Plans (NBSAPs) from member states of the Convention on Biological Diversity define the status of biodiversity in a country, the threats to biodiversity, and the strategies and actions to ensure conservation and sustainable use of biodiversity (CBD 2019). An aggregation of threats to biodiversity from sub-Saharan Africa NBSAPs would assist in understanding which threats are regional or sub-region and which are country-specific.

Threats to biodiversity include both direct and indirect threats. Direct threats are activities such as illegal hunting and agricultural expansion. Indirect threats are issue such as population growth and increasing per capita consumption of natural resources. As the causal linkages between indirect threats and biodiversity losses are often unclear or contested, analyses of biodiversity threats usually focus implicitly or explicitly on the direct threats to biodiversity (Carwardine et al. 2012; Dudgeon et al. 2006; Tolley et al. 2016). Direct threats can be defined as “the proximate human activities or processes that have caused, are causing, or may cause the destruction, degradation, and/or impairment of biodiversity” (Salafsky et al. 2008).

Here we systematically identify, triangulate, and rank the highest direct threats to biodiversity in sub-Saharan Africa and its sub-regions. To our knowledge, this is the first comprehensive and systematic assessment of direct biodiversity threats in sub-Saharan Africa and its sub-regions. Our aim is to respond to the research question: What are the highest direct threats to biodiversity in sub-Saharan Africa and its sub-regions? The primary target audience for this study are organizations seeking to develop an evidence-informed regional or sub-regional strategy for biodiversity conservation in sub-Saharan Africa. The secondary audience is conservation practitioners seeking to build partnerships or coalitions to address specific threats.

## 2 Methods

### 2.1 Threat classification

To classify and code the direct threats to biodiversity, we used the IUCN–Conservation Measures Partnership threats classification version 2.0 that comprises 43 different classification of direct threats to biodiversity (CMP 2016). While this classification system is contested given the complexity of threats to biodiversity (Balmford et al. 2009), the IUCN–CMP is the current standard used to classify and code biodiversity threats. This classification system also underpins the IUCN Red List threats classification system.

To develop a ranked list of direct threats to biodiversity conservation, we analyzed and combined threat assessments from three data sources. First, we created and implemented a formal knowledge elicitation process based on inputs from experts on sub-Saharan Africa biodiversity. Second, we used a global assessment of 33,044 IUCN Red-listed species that categorized the direct threats to each species (Maxwell et al. 2016) and from this dataset extracted the threats to sub-Saharan Africa threatened and near-threatened Red-listed species. Third, we used the National Biodiversity Strategy and Action Plans from sub-Saharan Africa countries (CBD 2019). Each of the methods for the three sources is detailed below. Averaging the threat rankings from the three sources provides our list of the highest threats to biodiversity conservation in sub-Saharan Africa and its sub-regions, along with an indication of sub-regional and source variability.

### 2.2 Expert knowledge

To elicit and assess expert knowledge about the direct threats to biodiversity, we used the Delphi method (Mukherjee et al. 2015). This method was developed in the 1950s (Dalkey and Helmer 1963) and refined over several decades (Linstone and Turoff 2002). Our Delphi follows Mukherjee et al. (2015): (i) preparation of first round of the questionnaire; (ii) selection and invitation of a panel of respondents; (iii) collection and analysis of the completed questionnaire for the first round; (iv) anonymous feedback on the responses gathered from all participants; (v) preparation and analysis of second round of questionnaire; and (vi) iteration.

A Delphi approach offers several benefits. By anonymizing expert inputs, a Delphi avoids social pressures such as groupthink, halo effect, egocentrism, and dominance by a few individuals (Mukherjee et al. 2015). The written feedback to the experts after each round makes the process retraceable and transparent. The sample size of the experts also matters less than the expertise and different perspectives of the respondents (Mukherjee et al. 2015).

A Delphi can suffer from several known issues (Groves et al. 2002; Martin et al. 2012; Mukherjee et al. 2015). First, there can be selection bias if participants are not representative of the population intended to be analyzed such as when a particular demographic group opts out of a survey. Attrition bias is also a risk if people who drop out of a survey are systematically different from those who stay. To assess selection and attrition bias, we collected data on respondents’ geographic areas of expertise and primary job in each round of the Delphi and analyzed the changes from round to round. Second, there can be linguistic uncertainties such as vagueness of terms. To minimize these, we used a vetted list of threats, pre-testing of the survey, and professional translation. Third, there may be nonresponse bias whereby the people who participate are systematically different from people who choose not to participate. To assess nonresponse bias, we conducted what is known as a wave analysis that compares responses from the initial wave of people taking the final round of the survey with the wave of people who completed the survey after the final reminder (a proxy for non-respondents) as per the guidance in Phillips et al. 2016.

For the Delphi, we developed an electronic survey that asked respondents to rank each of the 43 IUCN–CMP threats on a five-point Likert scale of very low, low, medium, high, and very high (1 to 5). We pre-tested the survey with 11 respondents and refined the wording of several questions. We then professionally translated the survey and cover email into French to widen the potential pool of respondents. Two native French speakers knowledgeable about biodiversity threats in Africa checked the quality of the translations.

The list of experts invited to participate was drawn from the co-authors’ sub-Saharan Africa contacts. We invited experts to participate who had two or more years of biodiversity conservation experience in two or more countries in sub-Saharan Africa. The experts themselves were also asked to nominate others who met the inclusion criteria. In total, we invited 217 experts to participate (17% from the co-authors’ organizations) and asked each participant to categorize their professional affiliation and position.

We developed a data analysis plan prior to the start of the Delphi and followed it during the analysis. The plan called for three rounds of expert elicitation. We set the threshold for expert consensus at 80% agreement (a common threshold) that a threat was high or very high. All responses were anonymous to avoid the halo effect.

In the first round, we invited respondents to suggest edits and additions to the threats. Based on input from respondents, we split hunting into legal and illegal hunting threats. As per the analysis plan, after the first round we dropped threats rated low or very low (1 or 2) by more than 50% of respondents. Thus, the second round began with 18 fewer threats. In the second round, we asked the experts who rated a threat very high to explain “why you rated this threat a 5 with words that might convince others.” After the second round, we kept those questions rated medium, high, or very high (3, 4 and 5) by 50% or more of respondents and those with at least one reason given for why it was a very high threat. The third round had eight fewer threats. In addition to ranking the overall threats for sub-Saharan Africa, we used data on the sub-regional expertise of each respondent to also rank the threat for the sub-regions of Central Africa, East Africa, Southern Africa, and West Africa.

### 2.3 Red-listed species

For the threats to IUCN Red-listed species, we used a global assessment of threats (Maxwell et al. 2016). This assessment included all assessed species that are categorized as threatened or near threatened. The assessment used the same IUCN–CMP threat classifications as our Delphi but an older version without multiple climate change categories. We modified this dataset to include only sub-Saharan Africa species following the IUCN list of sub-Saharan Africa countries and used the IUCN Red List API (IUCN 2017) and IUCN Red List Client package for R (Chamberlain 2017) to create a list of sub-Saharan Africa threatened and near-threatened species. After removing duplicates (i.e., single species present in multiple countries), we obtained a list of 21,121 unique species within sub-Saharan Africa. Using this list, we subset the Maxwell et al. (2016) threat data for species present in sub-Saharan Africa, resulting in a list of IUCN–CMP threats for 1,633 species. We then ranked the threats by frequency for sub-Saharan Africa and its four sub-regions.

### 2.4 NBSAPs

From the Convention on Biological Diversity website, we obtained the most recent NBSAPs (or the *Fifth National Report* if it was more recent than the NBSAP) for all sub-Saharan Africa countries except Gabon and Swaziland/eSwatini which do not have NBSAPs (*n* = 47). Each document includes a section on the direct threats to biodiversity in the country. The direct threats noted in the documents were entered into a database, checked by a second person for accuracy, and then mapped to the corresponding IUCN–CMP threat classification. Frequency counts for each threat were used to establish the NBSAP threat ranking for sub-Saharan Africa and its four sub-regions. The NBSAPs were largely from 2014, 2015 and 2016 (*n* = 41) with two older and four newer NBSAPs, so the time period was similar to the 2016 Delphi and 2016 Red-listed species data.

## 3 Results

### 3.1 Delphi sub-Saharan Africa threats

The Delphi had an average of 63 respondents across the three rounds, and the respondents had the varying expertise in Eastern Africa, Central Africa, Southern Africa, and Western Africa (Figure 1).

**Figure 1.**
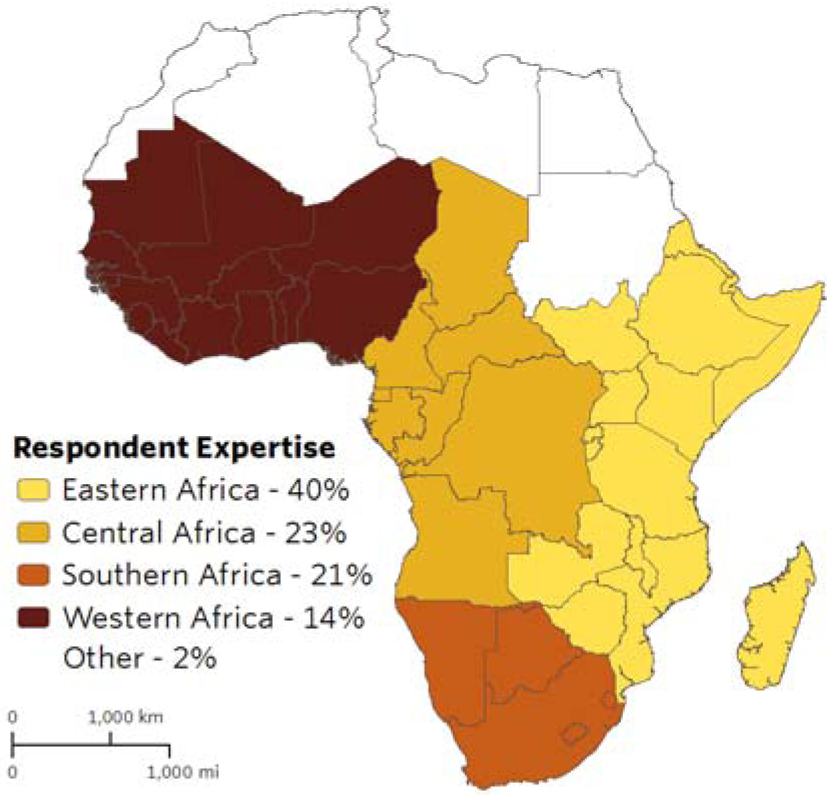
Map showing regional expertise of respondents.

Four direct threats to biodiversity in sub-Saharan Africa reached the threshold of 80% or greater expert consensus in the Delphi: crops, annual and perennial (non-timber) (87% consensus); fishing and harvesting aquatic resources (marine and freshwater) (86% consensus); illegal hunting and collecting terrestrial animals (84% consensus); and logging and wood harvesting (natural forests) (83% consensus). There was lower agreement on the threats ranked fifth, sixth, and seventh, with livestock ranching at 71% agreement, climate change/changes in precipitation at 68% agreement, and dams/water management at 53% agreement. Agreement dropped below the 50% level after the eighth ranked threat of mining/quarrying.

Regional expertise stayed largely consistent across the three Delphi rounds (8% variation) as did the percentage of respondents who worked for a non-governmental organization (4% variation). Given the similarity of respondent characteristics across Delphi rounds, attrition bias is unlikely to be a factor in the Delphi results.

Differences between those who work for a conservation NGO and those who do not were nominal; the same four threats were rated highest by both conservation NGO and non-NGO respondents, but the order changes slightly with illegal hunting the highest rated threat by non-NGO respondents.

In assessing possible nonresponse bias in round three (the round that generated the Delphi results), the wave analysis found a 0.1% difference in average threat ratings between the initial wave of people taking the round three survey and the wave of people taking the survey after the final reminder—a proxy for non-respondents. This nominal difference suggests nonresponse bias is unlikely to have influenced the Delphi results.

### 3.2 Red-listed sub-Saharan Africa threats

The top five threats to Red-listed species in sub-Saharan Africa are: crops, annual and perennial (non-timber); logging and wood harvesting (natural forests); housing and urban areas; fishing and harvesting aquatic resources (marine and freshwater); and invasive non-native/alien species. The frequency rankings for threats to Red-listed species in sub-Saharan Africa show several similarities and one difference compared to the Delphi. The highest ranked threat is the same for both (agriculture), and the rankings of logging and fishing are reversed but still in the top four. Yet hunting is ranked sixth in the Red-listed data and third in the Delphi. This is likely due to our splitting of the hunting threat in the Delphi into illegal and legal hunting threats which elevated illegal hunting in the rankings, while in the Red-listed data, legal and illegal hunting were combined.

### 3.3 NBSAP sub-Saharan Africa threats

Of the 47 NBSAP countries, 83% reported logging as a threat, 79% reported hunting was a threat, 77% reported agriculture as a threat, and 77% reported invasive species as a threat. Fishing, fire/fire suppression, and mining/quarrying were the three other threats with more than 50% of countries reporting them as direct threats to biodiversity conservation. Compared to the Delphi and Red-listed species top threats, logging moved to number one and agriculture dropped to number three.

When the rankings from the three data sources are averaged, there is a convergence around four threats: agriculture, logging/wood harvesting, fishing, and hunting. Several sources of pollution are also noteworthy for making the top ten threats. For climate change, only changes in precipitation/broad-scale hydrological regimes made the top ten (Table 2).

**Table 2.** Ten highest ranked threats to sub-Saharan African biodiversity.

| Threat | Average | Delphi Rank | Red-Listed Rank | NBSAP Rank |
| --- | --- | --- | --- | --- |
| Crops, annual and perennial (non-timber) | 1.7 | 1 | 1 | 3 |
| Logging and wood harvesting (natural forests) | 2.3 | 4 | 2 | 1 |
| Fishing and harvesting aquatic resources (marine and freshwater) | 3.7 | 2 | 4 | 5 |
| Hunting and trapping terrestrial animals | 3.7<sup>a</sup> | 3 | 6 | 2 |
| Livestock ranching and farming | 6.7 | 5 | 7 | 8 |
| Invasive non-native/alien species | 7.7 | 15 | 5 | 3<sup>b</sup> |
| Housing and urban areas | 9.3 | 16 | 3 | 9 |
| Pollution: industrial and military effluents | 9.5 | --<sup>b</sup> | 9 | 10 |
| Climate change - changes in precipitation and broad-scale hydrological regimes | 10.0 | 6 | 14 | 10<sup>a</sup> |
| Pollution: agricultural and forestry effluents | 10.5 | --<sup>b</sup> | 8 | 13 |

<sup>a</sup> Tied ranking
<sup>b</sup> Excluded after first round of Delphi because ranked by more than 50% of participants as a low threat

### 3.4 Sub-regional threats

The sub-regional rankings of threats exhibit more variation than the regional rankings. Overall, hunting is the top threat in Central Africa, agriculture is the top threat in East Africa, invasive species is the top threat in Southern Africa, and logging tied with agriculture as the highest threats in West Africa (Tables 4-7).

**Table 3.** Highest ranked threats to Central African biodiversity.

| Threat | Average | Delphi Rank | Red-Listed Rank | NBSAP Rank |
| --- | --- | --- | --- | --- |
| Hunting and trapping terrestrial animals | 1.7 | 1 | 3 | 1 |
| Logging and wood harvesting (natural forests) | 2.3 | 3 | 2 | 2 |
| Crops, annual and perennial (non-timber) | 3.0 | 3 | 1 | 5 |
| Fishing and harvesting aquatic resources | 3.0 | 2 | 4 | 3 |
| Pollution: agricultural and forestry effluents | 6.0 | --<sup>a</sup> | 7 | 5 |

<sup>a</sup> Excluded after first round of Delphi because ranked by more than 50% of participants as a low threat

**Table 4.** Highest ranked threats to East African biodiversity.

| Threat | Average | Delphi Rank | Red-Listed Rank | NBSAP Rank |
| --- | --- | --- | --- | --- |
| Crops, annual and perennial (non-timber) | 1.7 | 1 | 3 | 1 |
| Fishing and harvesting aquatic resources | 4.7 | 3 | 2 | 9 |
| Logging and wood harvesting (natural forests) | 5.0 | 5 | 7 | 3 |
| Pollution: agricultural and forestry effluents | 5.5 | --<sup>a</sup> | 4 | 7 |
| Recreational activities | 6.0 | --<sup>a</sup> | 6 | --<sup>b</sup> |

<sup>a</sup> Excluded after first round of Delphi because ranked by more than 50% of participants as a low threat
<sup>b</sup> Not mentioned in NBSAPs from the sub-region

**Table 5.** Highest ranked threats to Southern African biodiversity.

| Threat | Average | Delphi Rank | Red-Listed Rank | NBSAP Rank |
| --- | --- | --- | --- | --- |
| Invasive non-native/alien species | 2.3 | 4 | 2 | 1 |
| Crops, annual and perennial (non-timber) | 2.7 | 3 | 3 | 2 |
| Logging and wood harvesting (natural forests) | 3.7 | 2 | 5 | 4 |
| Fishing and harvesting aquatic resources | 5.0 | 7 | 1 | 7 |
| Climate change: changes in precipitation and broad-scale hydrological regimes | 7.5 | 4 | --<sup>a</sup> | 11 |

<sup>a</sup> Not included in Red-listed data because used version 1.1 IUCN-CMP threat classifications

**Table 6.** Highest ranked threats to West African biodiversity.

| Threat | Average | Delphi Rank | Red-Listed Rank | NBSAP Rank |
| --- | --- | --- | --- | --- |
| Crops, annual and perennial (non-timber) | 2.3 | 1<sup>a</sup> | 3 | 3 |
| Logging and wood harvesting (natural forests) | 2.3 | 1<sup>a</sup> | 5 | 1 |
| Fishing and harvesting aquatic resources | 3.3 | 4 | 1 | 5 |
| Hunting and trapping terrestrial animals | 3.3 | 1<sup>a</sup> | 7 | 2 |
| Invasive non-native/alien species | 7.3 | 16 | 2 | 4 |

<sup>a</sup>Tied

## 4 Discussion

Using triangulated data sources, we identified the highest threats for sub-Saharan Africa and each of its sub-regions. The top direct threats to biodiversity conservation in sub-Saharan Africa can be summarized as agriculture, logging/wood harvesting, fishing, and illegal hunting. The highest ranked threat in Central Africa is hunting. In East Africa, it is agriculture. In Southern Africa, it is invasive non-native/alien species, and in West Africa, agriculture and logging are tied as the highest threats. While none of these threats are new, the rankings highlight the primary direct threats for those organizations seeking to establish conservation priorities for sub-Saharan Africa or sub-regional strategies and conservation practitioners seeking to build coalitions to address biodiversity threats.

Known limitations of our results are the high percentage of participants in the Delphi who were from East Africa (40%) making selection bias a potential threat, and the ranking of threats in NBSAPs based on frequency, so threats identified in a number of smaller countries would rank higher than threats identified in one or two large countries even though the area impacted by the threat may be larger.

Our Africa-specific results are similar to a study using the same IUCN–CMP threat classifications as our study that ranked the threats to 1,961 protected areas across 149 countries (Schulze et al. 2018). Three of the four top threats in our results (hunting, logging, and fishing) fall within the top ten threats ranked by frequency in this study as well (Schulze et al. 2018).

The IPBES regional assessment report on biodiversity and ecosystem services for Africa includes a chapter on the direct and indirect drivers of change in biodiversity and nature contributions to people (IPBES 2018) and notes six “anthropogenic direct drivers”: land-use and land-cover change; deforestation; climate change; overexploitation; invasive alien species; and pollution. The four highest threats noted in our results are noted in the IPBES report as well, but the emphasis is different. The illegal wildlife trade is more prominent in the IPBES report, with the report’s qualitative assessment giving the illegal wildlife trade the highest level of agreement among the countries sampled.

Addressing threats to biodiversity is central to conservation success (Pressey et al. 2017), and several frameworks for systematic conservation planning include threats and their severity levels as key inputs (Ferrier and Drielsma 2010; Margules and Pressey 2000; Wilson et al. 2007). Other factors, however, are also relevant for systematic conservation planning including costs and benefits (Naidoo et al. 2006), spatial and temporal considerations (Pressey et al. 2007) and vulnerability (Wilson et al. 2005). Thus, identifying the main threats to biodiversity is an important step, but threats alone should not be the sole criterion for deciding on conservation investments.

Agriculture was a top three threat in all sub-regions of sub-Saharan Africa, and agricultural expansion is closely linked to population growth, consumption growth, and food security. Sub-Saharan Africa has the highest prevalence of hunger of any region globally, and the number of undernourished in the region increased from 2014 to 2017 (UN 2020). Expanding agricultural outputs in sub-Saharan Africa is critical for attaining the UN Sustainable Development Goal to end hunger, achieve food security and improved nutrition, and promote sustainable agriculture by 2030. A majority of the uncultivated arable land globally is found in sub-Saharan Africa (Roxburgh et al. 2010), and the region is a net importer of food (Van Ittersum et al. 2016). Moreover, closing the yield gap on existing cropland between current yields and potential yields will not be enough to make sub-Saharan Africa self-sufficient in cereal production by 2050 (Van Ittersum et al. 2016). Agricultural expansion will be needed, and sharing-sparing tradeoffs between agricultural expansion and wildlife habitat will continue (Phalan 2018). The severity of the tradeoffs, however, could be mitigated by environmental safeguards, channeling agricultural expansion to areas less important for biodiversity conservation, and accelerating the shift from “extensification,” whereby increased production comes from expanding into new land, to “intensification,” whereby yields are increased on existing farmland (Tilman et al. 2011). The long-term sustainability of intensive agriculture, however, is a contested question, and “sustainable intensification of agriculture” has been questioned because the sustainability of conventional intensification of agriculture is based on improved seed varieties, fertilizers, pesticides and herbicides (Kuyper and Struik 2014). Ecological intensification of agriculture, on the other hand, seeks to mimic natural ecosystems, incorporate traditional knowledge, and make use of comparative analysis of which cropping systems work well for which climate, soil, and socioeconomic conditions (Doré et al. 2011). Its utility, however, remains to be proven.

Within the logging/wood harvesting threat, logging to supply the wood products industry is different than wood harvesting to supply household energy needs. The former is tied to global markets and trends while the latter is more localized. A majority of the wood harvesting in sub-Saharan Africa is for households consumption, and per capita wood consumption is two to three times higher than elsewhere in the world with an increasing trend line (FAO 2017). Giving local people secure tenure or management and benefit rights to local forest makes them more likely to establish and follow usage rules and monitor the compliance of others (Ostrom and Nagendra 2006). Providing local performance payments to keep forests intact by developing avoided deforestation carbon projects also shows promise in sub-Saharan Africa (Anderson et al. 2012).

Demand for food fish in sub-Saharan Africa is projected to grow by 30% between 2010 and 2030 (Msangi et al. 2013), and illegal, unreported and unregulated fishing (IUU) are known issues in sub-Saharan Africa (Braulik et al. 2017; Daniels et al. 2016). Commercial IUU fishing is particularly acute in West Africa, and a number of suggestions for improvements have been proposed such as blacklisting IUU vessels and creating a global database and tracking system (Daniels et al. 2016). Among small-scale fishers in both freshwater and marine environments, creating no-take zones/fish regeneration areas and locally managed marine areas (LMMAs) have worked elsewhere and show promise in sub-Saharan Africa (Rocliffe et al. 2014; Samoilys and Obura 2011).

Wildlife for food is a critical part of indigenous peoples’ diets in parts of sub-Saharan Africa. It substitutes for a lack of sufficient and affordable protein available in rural areas and fills a social/cultural/gastronomic role in urban areas (Wilkie et al. 2016). Hunting also adds to household income. Known tools to reduce illegal hunting include devolution of wildlife management authority to communities with legitimate claims to land (Cooney et al. 2017; Roe et al. 2015), anti-poaching patrols (Hilborn et al. 2006), demand reduction (Vigne and Martin 2017), protein substitution (Besbes et al. 2012), technology such as drones to reduce human-wildlife conflict and detect poachers (Hahn et al. 2017; Mulero-Pázmány et al. 2014), and community members trained to avert human-wildlife conflicts (Chang’a et al. 2016). Despite these known tools, investment at the scale necessary to address hunting threats to wildlife is currently lacking.

Finally, the complex social-ecological systems in which most biodiversity conservation happens mean that when one factor changes, another factor can change as well (Game et al. 2014). Addressing logging or IUU fishing, for example, can have knock-on effects on agricultural expansion and illegal hunting. Thus, addressing one threat in isolation may have marginal benefits to conservation (Wilson et al. 2006). Overall, if large conservation funding organizations target a high percentage of conservation investments towards the highest threats to biodiversity and these organizations encourage conservation practitioners to address the highest threats in partnership with others, then this would reduce biodiversity losses in sub-Saharan Africa.

## Acknowledgements

We thank all the participants who contributed their time and knowledge to the three Delphi rounds. We thank Sean Maxwell at University of Queensland for sharing his threats and IUCN Red-listed species dataset and for comments on the draft manuscript, and we thank Tracy Baker at The Nature Conservancy for creating the map.

